# Alcohol Cues Invigorate Aversion-Resistant Alcohol Drinking

**DOI:** 10.64898/2026.07.14.738545

**Authors:** Meredith R. Bauer, Jocelyn M. Richard

**Affiliations:** University of Minnesota, Department of Neuroscience

**Keywords:** Aversion-resistant, quinine, cues, alcohol, motivation

## Abstract

**Background:** Alcohol use disorder is characterized by continued alcohol use despite negative consequences, also known as aversion-resistant drinking. Alcohol related cues can invigorate motivation to seek and consume alcohol. It is currently unknown whether alcohol related cues can invigorate drinking despite negative consequences.

**Materials and Methods:** Long-Evans rats were trained in a discriminative stimulus (DS) task with cues predicting the availability of an alcohol reward. They were then tested in the task for aversion-resistant drinking by measuring consumption of alcohol adulterated with increasing concentrations of quinine. As a control for an environment free of explicit reward-related cues, rats were also tested for aversion-resistant drinking in the home cage.

**Results:** We found that rats displayed robust aversion-resistant drinking in the DS task. When we compared alcohol consumption in the task with home cage consumption, we found that rats were more aversion-resistant in the task than in the home cage. We also found that individual differences in aversion-resistant drinking were correlated within behavioral context (i.e. home cage or DS task) but not between the home cage and the DS task.

**Conclusions:** We found that aversion-resistant drinking is invigorated during cue-induced alcohol seeking relative to free drinking without explicit alcohol cues. Home cage quinine-sensitivity is unrelated to quinine-sensitivity in the presence of cues. This suggests that cues motivate drinking despite negative consequences in a way that is unique from aversion-resistance driven by drinking history. While many behavioral measures use cues and ultimately test aversion-resistant drinking, this is the first explicit test of cue-evoked aversion-resistant drinking.

## INTRODUCTION

Alcohol use disorder (AUD) is a chronic relapsing disease characterized by continued alcohol use despite negative consequences (American Psychological Association, 2022). Stark real-world examples of drinking despite negative consequences include drinking despite health issues such as liver cirrhosis and non-beverage alcohol consumption whereby an individual might consume an alcohol containing solution (e.g., mouthwash, cologne, antifreeze) despite it being highly aversive in taste and potentially containing poisonous chemicals like methanol. This behavior is frequently modeled in rodents by adding a bitter tasting substance, quinine, to alcohol solutions. Under normal conditions, rats will drink alcohol freely but suppress their consumption of alcohol when it is adulterated with quinine. However, under some conditions the drinking persists despite the bitter taste, effectively modeling drinking despite negative consequences. For example, a history of alcohol drinking (Bauer et al., 2021; Hopf et al., 2010; Lesscher et al., 2010) and genetics (Houck et al., 2019; Timme et al., 2020) can promote quinine-adulterated alcohol consumption. This phenomenon is referred to as aversion-resistant alcohol drinking. The underlying cause of aversion-resistant drinking is not fully known.

Another important factor that may influence aversion-resistant drinking is the presence of alcohol-related sensory stimuli, or cues. Cues that predict the availability or delivery of alcohol and other drugs are thought to promote addictive behavior via their incentive properties, which can produce complex motivational and emotional states that may lead to “compulsive” aversion-resistant alcohol seeking and consumption (Robinson & Berridge, 1993; 2001; 2025; Berridge & Robinson, 2016). Because alcohol is legal, socially acceptable to consume, and commonly used in society, avoiding environments with alcohol and alcohol-related cues is particularly difficult. Thus, a Gordian knot for recovery from and prevention of AUD is navigating environments fraught with alcohol-related cues. Understanding the relationship between alcohol-related cues and aversion-resistant drinking may aid in the prevention and treatment of AUD. However, while many assessments of aversion-resistant drinking involved implicit or explicit cues, the influence of these cues on aversion-resistant drinking has not been directly tested.

Thus, our goal was to causally test whether alcohol-related cues invigorate aversion-resistant drinking. To do so we used a discriminative stimulus (DS) task where one cue (the DS) predicts the availability of an alcohol reward and another cue (the NS) predicts nothing. We compared consumption of quinine-adulterated alcohol in the DS task to consumption in an environment that is relatively free of explicit alcohol cues. We found that animals continued to consume alcohol that had been adulterated with quinine to a greater extent in the DS task than in the home cage. This suggests that environments with alcohol-related cues can invigorate aversion-resistant drinking. Together, these data shed light onto the powerful role that alcohol-related cues play in invigorating aversion-resistant drinking and provide a model in which the neurobiological mechanisms of cue-elicited aversion-resistant drinking can be tested.

## MATERIALS AND METHODS

### Subjects

Adult male (n = 5) and female (n = 5) Long-Evans rats (Envigo) were individually housed with ad libitum access to food and water (Teklad Global 18% Protein Rodent Diet) and were maintained on a 14h-10h light-dark cycle. All experimental procedures were approved by the Institutional Animal Care and Use Committee at the University of Minnesota and carried out in accordance with the guide for the Care and Use of Laboratory Animals (NIH).

### Alcohol Preexposure

The purpose of the alcohol preexposure is to give rats experience with the pharmacological effects of alcohol and to increase the motivation for rats to respond in the task for an alcohol reward. All rats were given four weeks of intermittent access to 15% alcohol in their home cages. Alcohol bottles were added on Sunday, removed 24-hours later on Monday, and this pattern repeated through Friday. Rats had 48-hours without alcohol access from Friday bottles off to Sunday bottles on. This repeated for four weeks. Rats had access to food and water continuously throughout alcohol pre-exposure.

### Discriminative Stimulus (DS) Task

The DS task is an instrumental cue-based task where rats have the opportunity to earn 15% alcohol if they respond during a DS auditory cue, but not if they respond during an NS control cue. Training occurred in operant chambers equipped with speakers, lights, tone generators, and reward ports (Med Associates Inc., Fairfax, VT). Both auditory cues were set at 60-65 decibels. Rats first underwent 3, 90-minute sessions of magazine training where they had the opportunity to receive up to 30, 0.1 ml, 15% alcohol rewards to acclimate them to the operant chambers and reward ports. Rats then began training in the DS task across daily (M-F) one-hour sessions for a 0.06 ml alcohol reward. First, rats underwent 14 days of training where the only cue presented was the DS. DS cues were presented in decreasing lengths of time (60s, 30s, 20s, 10s) across days based on task performance. When rats responded during 60% of the trials in a session, they were moved to the next stage of training until they reached the 10s DS length. On session 15, a 10-second neutral stimulus (NS) was introduced in the task. Responding during the NS resulted in nothing. Rats trained with both the DS and NS cues for 8 sessions. The DS was a white noise, and the NS was a tone. The DS and the NS were presented pseudorandomly on a variable interval schedule with an average interval of 30 seconds, and each cue-type lasted up to 10s.

After a total of 21 days of training in stages and with both cue types, aversion-resistant drinking was tested (detailed description below). Following initial aversion-resistant drinking testing the reward volume was increased to allow rats the opportunity to titrate their consumption levels to their preference and to closer align consumption volumes in the DS task to the home cage. The volume of liquid rats were given per each reward was increased from 0.06 to 0.18 ml. After three days female rats significantly reduced their responding and consumption, so they were given 0.12 ml/reward which restored response rates. Males maintained 0.18 ml/reward as their responding and consumption was stable. Aversion-resistant drinking was tested again once responding at the new reward volume stabilized.

### Quinine-Adulterated Alcohol

To test for aversion-resistant drinking rats were given alcohol solutions adulterated with increasing concentrations of quinine. Consumption was compared with 0 mg/L (15% alcohol-only) consumption levels. Concentrations of quinine-alcohol were 30, 60, 90, and 180 mg/L. Solutions were presented in order from lowest to highest concentration across the sessions for each week. Sensitivity to quinine-alcohol was tested in the DS task at regular and high volumes and in the home cage.

### Home Cage Drinking

Rats were tested for aversion-resistant drinking in the home cage following testing in the DS task. Rats had access to a 15% alcohol bottle for one hour. Bottles were weighed immediately prior to being placed on the cage and immediately following bottle removal. Rats did not have access to water during the home cage sessions. Rats were given 6 sessions of home cage drinking prior to the addition of quinine to the alcohol bottles across four concentrations (30, 60, 90, and 180 mg/L).

### Statistical Analyses

Data analyses were conducted using MATLAB (Mathworks) and R (The R Project). Analyses ran were linear mixed-effects models that used subject as a random intercept to account for variability between subjects. This was followed by ANOVA. Post-hoc Dunnett’s or Student’s t-test were used with Bonferroni correction to assess significant interactions following Shapiro-Wilk test for normality. Effect sizes were calculated using generalized eta squared and were interpreted using Cohen’s D (Cohen, 1992). Pearson’s correlation was used to assess relationships between DS task and home cage quinine-sensitivity. Statistical significance was set at p < 0.05 for all tests and trends up to p = 0.1 are reported in the main text. All non-significant results are listed in Supplementary Tables 1-3.

## RESULTS

### Rats learn the DS task with an alcohol reward

Prior to training in the DS task, rats underwent four weeks of alcohol preexposure in an intermittent access two-bottle choice procedure with 15% alcohol. Alcohol consumption did not differ across day, sex, or interact between day and sex (Fig. 1B; p’s > 0.10). During training in the DS task rats increased their port entry probability across sessions, suggesting that rats learned that port entries during the DS result in a reward (Fig. 1C, main effect of day: F(1, 210) = 313.22, p < 0.0001; no significant effect of sex or interaction of sex and day, p’s > 0.10). When we compared port entries during the DS and NS, starting when the NS was introduced during session 15, we found significant discrimination between the DS and NS, which improved across days (Fig. 1C, main effect of cue type: F(1,150) = 44.38, p < 0.0001, η² = 0.23; main effect of day: F(1,150) = 12.84, p < 0.0001, η² = 0.08; interaction between day and cue type: F(1,150) = 8.08, p = 0.005, η² = 0.05). While we observed an impact of sex (main effect of sex: F(1,15.97) = 4.62, p = 0.047, η² = 0.22; interaction between sex and cue type: F(1,150) = 3.01, p = 0.08, η² = 0.02), males and females reached similar levels of discrimination by the end of training. Rats also increased in the number of rewards received over the course of training which further suggests that rats understood the task (Fig. 1D, main effect of day: F(1, 210) = 313.91, p < 0.001, η² = 0.60, no significant effect of sex or interaction of day and sex, p’s > 0.10). Alcohol consumption varied by day and sex (Fig. 1E, main effect of day: F(1,210) = 9.36, p = 0.0025, η² = 0.04; no significant effect of sex (p’s > 0.10), interaction of sex and day: F(1,210) = 17.85, p < 0.001, η² = 0.08), with females consuming significantly more grams per kilogram alcohol at the end of training. This is due in part to the limited amount of alcohol available in the task and differences in weight between males and females. Together, these data indicate that rats successfully learned the DS task.

**Figure 1.**
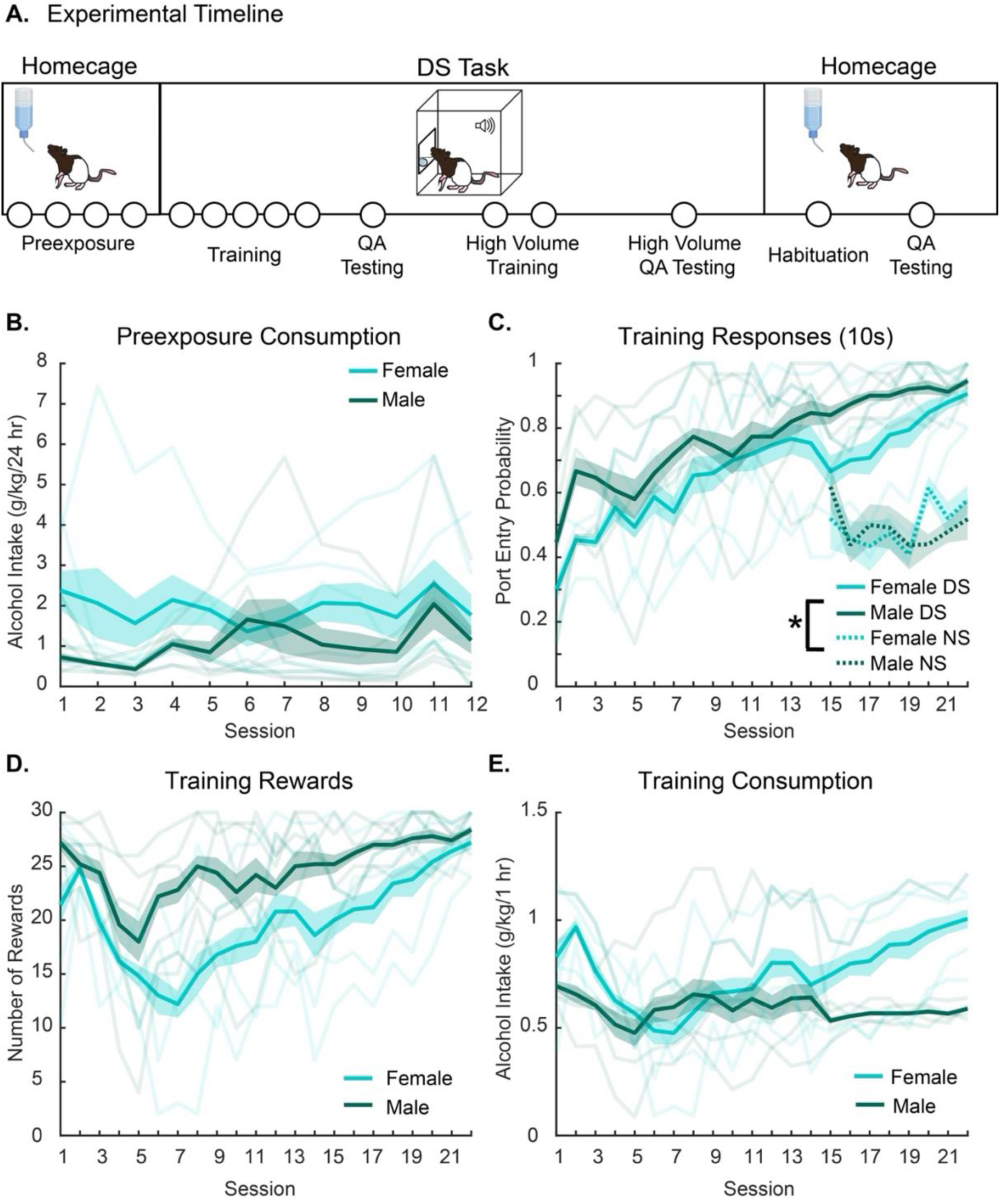
Rats learn the DS task with an alcohol reward. **A.** Experimental timeline, each dot represents approximately 5 days. **B.** Rats consume alcohol during alcohol preexposure. **C.** Rats increase their responding for the DS over time and discriminate between the DS and NS. **D.** Rats receive most of the rewards available to them through responding on the DS. **E.** Rats consume alcohol during the one-hour DS task sessions. Asterisks (*) indicate a significant difference.

### Rats demonstrate aversion-resistant drinking in the DS Task

Rats were then tested for aversion-resistant drinking in the DS task. We added increasing concentrations of quinine (30, 60, 90, and 180 mg/L) to the alcohol reward solutions and measured performance and consumption in the DS task. DS responses were unaffected by quinine adulteration or sex (Fig. 2A, p’s > 0.10). To account for individual differences in initial seeking and consumption, we assessed change scores from the baseline (five-day average prior to quinine testing) behavior. Percent change in DS responses was unaffected by quinine adulteration or sex (Fig. 2B, p’s > 0.10). In contrast, quinine adulteration decreased NS responding (Fig. 2C, main effect of quinine concentration: F(5,50) = 18.68, p < 0.0001, η² = 0.65; no significant effect of sex or interaction of quinine concentration and sex, p’s > 0.10). Post-hoc Bonferroni corrected Dunnet’s t-test determined that both 90 (*diff* = −.23, 95% CI [−.39, −.08], p < 0.001) and 180 mg/L (*diff* = −.24, CI [−.39, −.08], p < 0.001) quinine adulteration significantly reduced NS responses compared with baseline responding. Percent change in NS responses was also reduced by quinine concentration (Fig 2D, main effect of quinine concentration: F(1,50) = 17.59, p < 0.0001, η² = 0.26; no significant effect of sex or interaction of sex and quinine concentration, p’s < 0.05). Post-hoc Bonferroni corrected Dunnet’s t-test determined that both 90 mg/l quinine adulteration significantly reduced NS responses compared with responding for alcohol adulterated with 0 mg/L quinine (*diff* = −31.79, 95% CI [−61.57, −2.02], p = 0.03) and 180 mg/L (*diff* = −34.26, CI [−64.03, −4.48], p = 0.02). One hypothesis as to why we see this effect in NS responding is that rats might improve in their task performance over the course of the week as the training schedule is Monday through Friday. We assessed whether NS responding decreased from Monday to Friday across our training data and found that NS responding was not affected by day of the week when we isolated the final 5 days (Monday – Friday) of training (Fig. 1C, main effect of day: F(1,40) = 3.91, p = 0.06, η² = 0.09; no significant effect of sex or interaction of day and sex, p’s > 0.10). Since the p-value of the main effect of day trended towards significance, though despite a small effect size, we ran post-hoc Dunnet’s t-test with Bonferroni corrections which further confirmed that NS responding did not differ on Monday compared with the latter days of the week (all p’s > 0.10). Thus, the decrease in NS responding across increasing concentrations of quinine is likely related to the increasingly bitter alcohol solution constraining alcohol seeking outside of the DS period. We next analyzed port entry latency following cue onset for the DS and NS. Latency to respond to the DS was not affected by sex or quinine concentration (p’s > 0.10, Fig. 2E). Latency to respond to the NS was not affected by sex or quinine concentration (p’s > 0.10), though there was a significant interaction of sex and quinine concentration (Fig. 2F, F(5, 50) = 2.41, p = 0.049, η² = 0.19). Post-hoc tests determined there were no significant pairwise differences that explained this interaction (p’s > 0.10). Percent change in latency to respond are shown in Supplementary Figure 1. Finally, we assessed whether quinine concentration altered consumption of the alcohol during the task and found that quinine adulteration did not alter consumption (Fig. 2E, main effect of sex: F(1,10) = 31.13, p < 0.0001, η² = 0.76, no significant effect of quinine concentration or interaction of sex and quinine concentration, p’s > 0.10), or percent change in consumption (Fig. 2F, no significant effect of sex, quinine concentration, or interaction of sex and quinine concentration, p’s > 0.10). Thus, rats showed aversion-resistant drinking in the DS task, even at relatively high concentrations of quinine.

**Figure 2.**
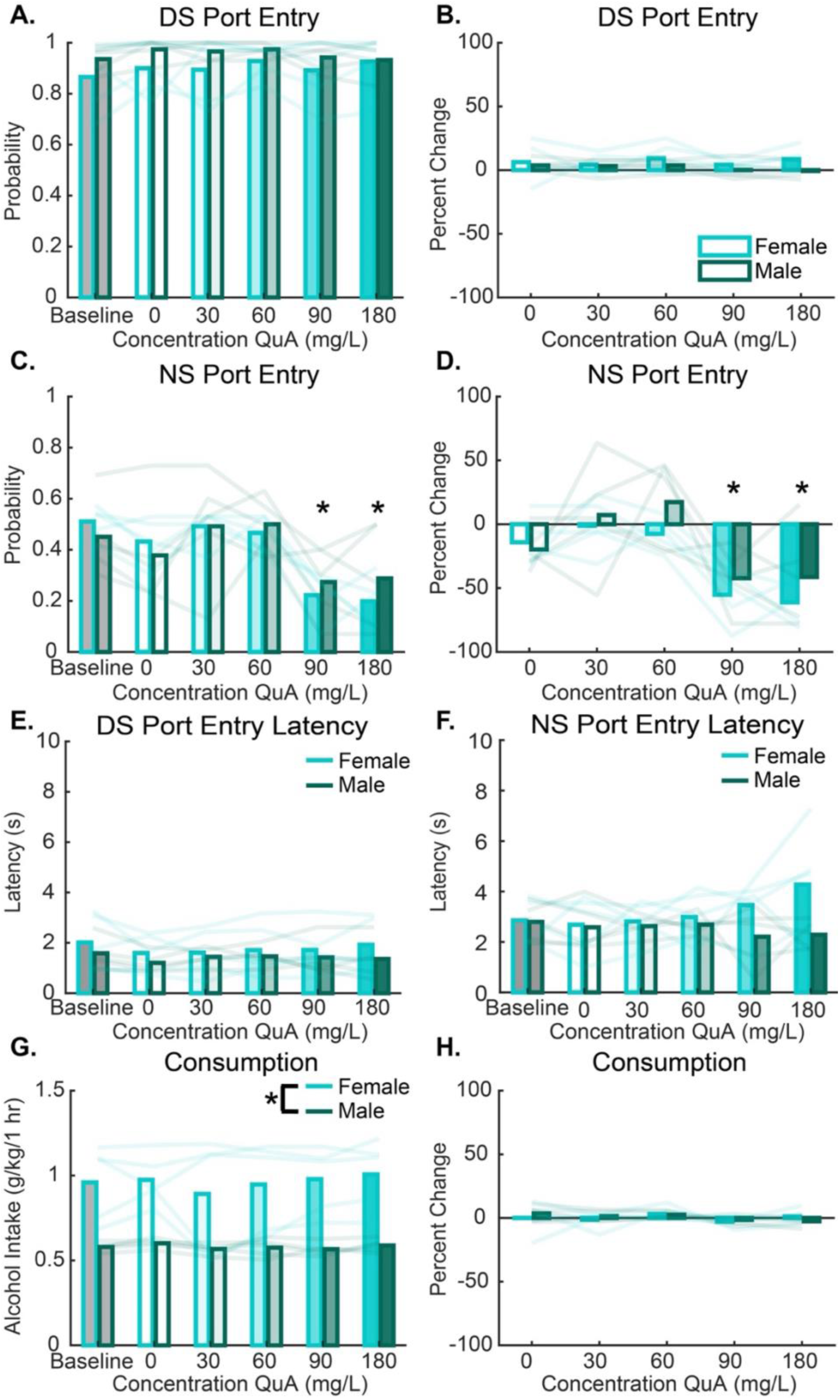
Rats demonstrate aversion-resistant drinking in the DS task. **A.** DS responses were not affected by increasingly aversive quinine-alcohol solutions. **B.** DS responses as a percent change from baseline were not affected by increasingly aversive quinine-alcohol. **C.** NS responses were significantly reduced at 90 and 180 mg/L quinine-alcohol as compared with baseline. **D.** NS responses as a percent change from baseline were significantly reduced at 90 and 180 mg/L quinine-alcohol compared with 0 mg/L quinine-alcohol. **E.** Latency to respond to the DS was not affected by quinine concentration. **F**. Latency to respond to the DS was not affected by quinine concentration. **G.** Reward consumption was not affected by increasingly aversive quinine-alcohol solutions. Females had higher g/kg consumption than males. **H.** Reward consumption as a percent change from baseline was not affected by increasingly aversive quinine-alcohol solutions. Asterisks (*) indicate a significant difference.

### Aversion-resistant drinking is heterogeneous at higher reward volumes

We hypothesized that cues evoke aversion-resistant drinking. To causally test this, we sought to compare DS aversion-resistant drinking with quinine intake in a cue-free, free-access drinking context in the home cage. However, a large difference between DS task consumption and home cage drinking is that rats can titrate their consumption levels in the home cage. To address this, we increased the reward volume in the DS task so that rats received more alcohol and would be able to titrate their consumption levels. Additionally, the increased reward volume would also increase the amount of time the rat interacted with the aversive solution. We hypothesized that this would increase the opportunity for the rats to experience the aversive properties of the solution to a greater extent than with the lower reward volume.

Following aversion-resistant drinking testing with the normal reward volume, we shifted the rats to a higher volume per reward where male and female rats went from receiving a 0.06 ml reward up to 30 times in a session to receiving a 0.18 ml reward up to 30 times in a session. Rats continued to discriminate between the DS and the NS across nine high volume training sessions (Fig. 3A, main effect of day: F(1,170) = 39.79, p < 0.001, η² = 0.19; main effect of cue type: F(1,170) = 184.28, p < 0.001, η² = 0.52; interaction of sex and day: F(1,170) = 5.02, p = 0.03 η² = 0.03; interaction of day and cue type: F(1,170) = 3.76, p = 0.05, η² = 0.02; interaction of sex, day, and cue type: F(1,170) = 2.93, p = 0.09, η² = 0.02, and no significant effect of sex, p > 0.10). However, despite male and female rats continued responding in the task at the higher reward volumes, female rats reduced their reward consumption and port entry probability across days (Fig. 3B, main effect of sex: F(1,49.2) = 75.9, p < 0.001, η² = 0.61; main effect of day: F(1,80) = 71.89, p < 0.0001, η² = 0.47; interaction of sex and day: F(1,80) = 71.51, p < 0.001, η² = 0.47). To stabilize port entry probability and alcohol consumption, we modified the reward volume for females from 0.18 ml/reward to 0.12 ml/reward to ensure they continued to engage in the task. This volume shift occurred on day 6 and was maintained for the rest of the experiment. Consumption remained steady for both males and females in the last four days of high-volume training (i.e., days 6-9) with the volume adjustment (Fig. 3B, main effect of day: F(1,30) = 3.69, p = 0.06, η² = 0.11; no significant effect of sex or interaction of sex and day, p’s > 0.10).

**Figure 3.**
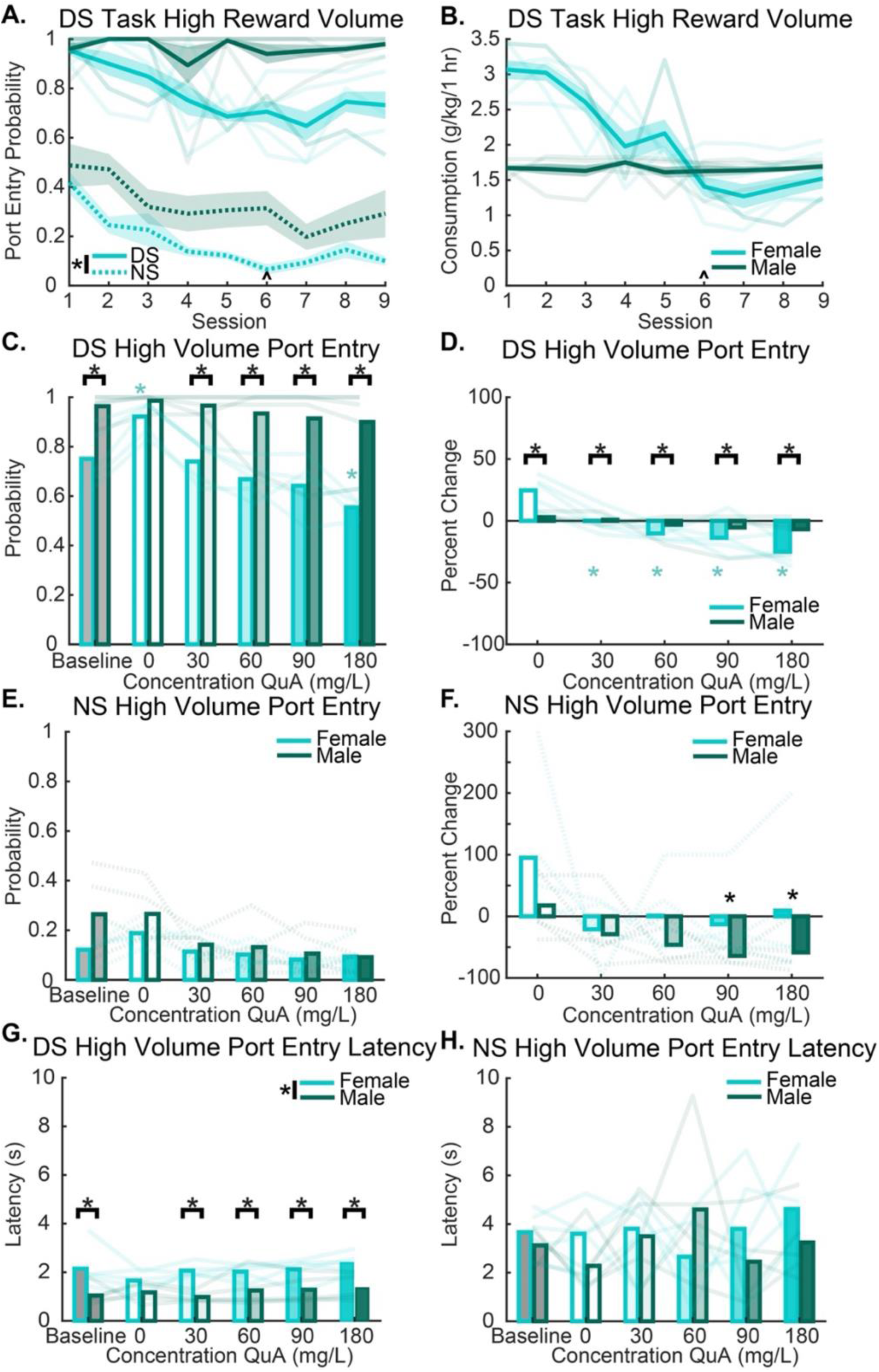
Aversion-resistant drinking is heterogeneous at higher reward volumes. **A.** Reward volume was increased (to 0.18 ml) and rats trained with the higher volume across 9 sessions with the exception that reward volume was reduced at session 6 for females (to 0.12 ml/reward), but not males. The day that the change in reward volume occurred is indicated on the plot with a caret (^). Rats continued to respond during the DS and discriminated between the DS and NS. **B.** Rats consumed alcohol across the 9 sessions. Consumption levels reduced during the 0.18 ml/reward access (sessions 1-5) and stabilized once females were moved to 0.12 ml/reward (sessions 6-9). The day that the change in reward volume occurred is indicated on the plot with a caret (^). **C.** As compared with the averaged baseline days (Baseline), DS responses were significantly reduced at 180 mg/L compared with baseline for female rats only. Female rats responded significantly more on the 0 mg/L quinine-alcohol day compared with baseline. Further, males responded significantly more than females at baseline, 30, 60, 90, and 180 mg/L days. **D.** DS responses as percent change from baseline demonstrated that female rats significantly reduced their DS responses relative to baseline at all quinine-alcohol concentrations. Additionally, females responded significantly more than males at 0 mg/L, but significantly less than males at all concentrations of quinine-alcohol. **E.** NS responses were not affected by quinine-alcohol concentration as compared with baseline. **F.** NS responses as percent change from baseline were significantly reduced at 90 and 180 mg/L quinine-alcohol. **G.** Latency to respond to the DS was significantly higher for females versus males at baseline, 30, 60, 90, and 180 mg/L. **H**. Latency to respond to the NS was not affected by quinine concentration or sex. Color coded asterisks (*) indicate a significant difference within sex, black asterisks with a bar above them indicate a significant difference between males and females.

Rats were then tested for aversion-resistant drinking in the DS task under identical procedures as before with the only exception being the higher reward volume. Under these conditions, DS responding was reduced by the quinine adulteration (Fig 3C, main effect of quinine adulteration concentration: F(5,50) = 19.35, p < 0.0001, η² = 0.66). We also observed greater responding in males overall (main effect of sex: F(1,10) = 23.71, p < 0.0001, η² = 0.70) which depended on quinine concentration (interaction of sex and quinine concentration: F(5,50) = 6.97, p < 0.0001, η² = 0.41). Post-hoc Dunnet’s test in females revealed a significant difference between baseline and 0 mg/l quinine (*diff* = 0.17, 95% CI [0.02, 0.32], p = 0.02) and between baseline and 180 mg/l quinine (*diff* = −0.20, 95% CI [−0.35, 0.05], p = 0.007) suggesting maintained task engagement in females up until 180. Post-hoc Dunnet’s t-test in males revealed no differences between baseline and any quinine concentration suggesting unwavering responding despite the addition of aversive taste. Comparison of males versus females at each concentration using post-hoc Welch’s Two Sample t-test revealed that males responded more than females during the baseline days (*t*(5.87) = −3.73, p = 0.01), 30 mg/L quinine (*t*(5.37) = −5.59, p = 0.002), 60 mg/l quinine (t(−3.93) = 6.35, p-value = 0.007), 90 mg/L quinine (*t*(6.85) = −2.90, p = 0.02), 180 mg/L (*t*(4.98) = −4.67, p = 0.006), but there was no sex difference at 0 mg/L quinine (*t*(5.66) = −1.93, p = 0.11). These data suggest that males are less affected by quinine adulteration in their responding and both male and female rats demonstrate aversion-resistant responding.

To account for individual differences, we assessed change scores from the baseline behavior days. We found that DS responses reduced across quinine concentration and this effect depended on sex (Fig. 3D, main effect of quinine concentration: F(4,40) = 23.08, p < 0.0001, η² = 0.70; interaction of sex and quinine concentration: F(4,40) = 9.95, p < 0.0001, η² = 0.50; no significant effect of sex, p > 0.10). Post-hoc Dunn’s test with Bonferroni correction comparing 0 mg/L to the quinine concentrations revealed a significant difference compared with 60 (z = −20.40, p = 0.01), 90 (z = −20.35, p = 0.02), 180 (z = −27.00, p < 0.001), but not 30 (z = −12.00, p = 0.63). Post-hoc Dunnet’s t-test with Bonferroni correction for each sex demonstrated that this effect was driven by females (Females 0 mg/L vs: 30 mg/L (*diff* = 24.73, 95% CI [−43.41, −6.047], p = 0.008); 60 mg/L (*diff* = −34.84, 95% CI [−53.52, −16.16], p = 0.0003); 90 mg/L (*diff* = −38.33, 95% CI [−57.01, −19.65], p < 0.0001); 180 mg/L (*diff* = −49.54, [−68.22, −30.86], p < 0.0001), as there were no significant differences between 0 mg/L and any quinine concentration in males (all p’s > 0.10). Post-hoc Welch’s test comparing males versus females at each concentration revealed that females responded more than males at 0 mg/L: (*t*(4.51) = 3.75, p = 0.02), and responded less than males at all other concentrations of quinine-alcohol; 30 mg/L: (*t*(−5.59) = 5.37, p = 0.002), 60 mg/L: (*t*(6.35) = −3.93, p = 0.007), 90 mg/L: (*t*(6.89) = −2.90, p = 0.02), and 180 mg/L: (*t*(4.98) = −4.67, p = 0.006).

Similar to our finding in the low-volume version of the DS task, NS responses were reduced by quinine adulteration (Fig 3E, main effect of quinine concentration: F(5,50) = 9.8, p < 0.0001, η² = 0.50; no significant effect of sex (p > 0.10); interaction of sex and quinine concentration: F(5,50) = 2.18, p = 0.07, η² = 0.18), though this effect was not explained by any individual quinine concentration (Post-hoc Bonferroni corrected t-tests comparing baseline NS responses with all other concentrations (p’s > 0.10)). Percent change in NS responding significantly reduced across quinine concentration (Fig. 3F, main effect on quinine adulteration concentration: F(4,40) = 5.76, p < 0.0001, η² = 0.37; main effect of sex: F(1,10) = 3.82, p = 0.08, η² = 0.276; no significant interaction of sex and quinine concentration (p > 0.10)). Post-hoc Dunn’s tests with Bonferroni corrections comparing 0 to quinine adulteration each quinine concentration revealed a significant difference compared with 90 mg/L (z = −20.10, p = 0.02) and with 180 mg/L (z = −19.85, p = 0.02), but not 30 mg/L or 60 mg/L (p’s > 0.10). Next, we looked at port entry latency following the cue for the DS and the NS. DS port entry latency was significantly affected by quinine concentration, sex, and interacted by sex and quinine concentration (Fig. 3G: main effect of quinine concentration, F(5, 48.18) = 3.37, p = 0.01, η² = 0.259); main effect of sex, F(1, 10.09) = 18.64, p = 0.0015, η² = 0.649; and interaction of quinine concentration and sex: F(5, 48.18) = 2.88, p = 0.0237, η² = 0.23). Post-hoc t-tests determined this interaction was driven by females responding slower than males at 30 mg/L (p = 0.0001), 90 mg/L (p = 0.042), and 180 mg/L (0.0084). NS response latencies were unaffected by sex, quinine concentration, or interaction of sex and quinine concentration (p’s > 0.10). Percent change in latency is shown in Supplemental Figure 1. These data further suggest that rats constrain their responding for rewards to the DS during aversive quinine sessions.

### Rats demonstrate greater aversion-resistant drinking in the DS task compared with home cage drinking

We then assessed quinine-alcohol consumption in the higher reward volume. Quinine adulteration significantly reduced consumption (Fig 4A, main effect of quinine concentration: F(5,50) = 19.14, p < 0.0001, η² = 0.66; no significant effect of sex, (p > 0.10); interaction of sex and quinine concentration: F(5,50) = 3.04, p = 0.02, η² = 0.23). Post-hoc Bonferroni corrected Dunnet’s t-test determined that 180 mg/L quinine significantly reduced consumption compared with baseline drinking (*diff* = −0.55, 95% CI [−0.93, −0.18], p = 0.002). This effect was driven by females as post-hoc Dunnet’s t-test with Bonferroni correction in females only revealed that 180 mg/L reduced drinking compared with baseline (*diff* = −.68, 95% CI [−1.16, −0.19], p = 0.004). Post-hoc Dunnet’s t-test in males revealed no significant differences compared with baseline (all p’s > 0.10). Males and females did not differ significantly in their consumption at any quinine concentration (p’s > 0.10).

**Figure 4.**
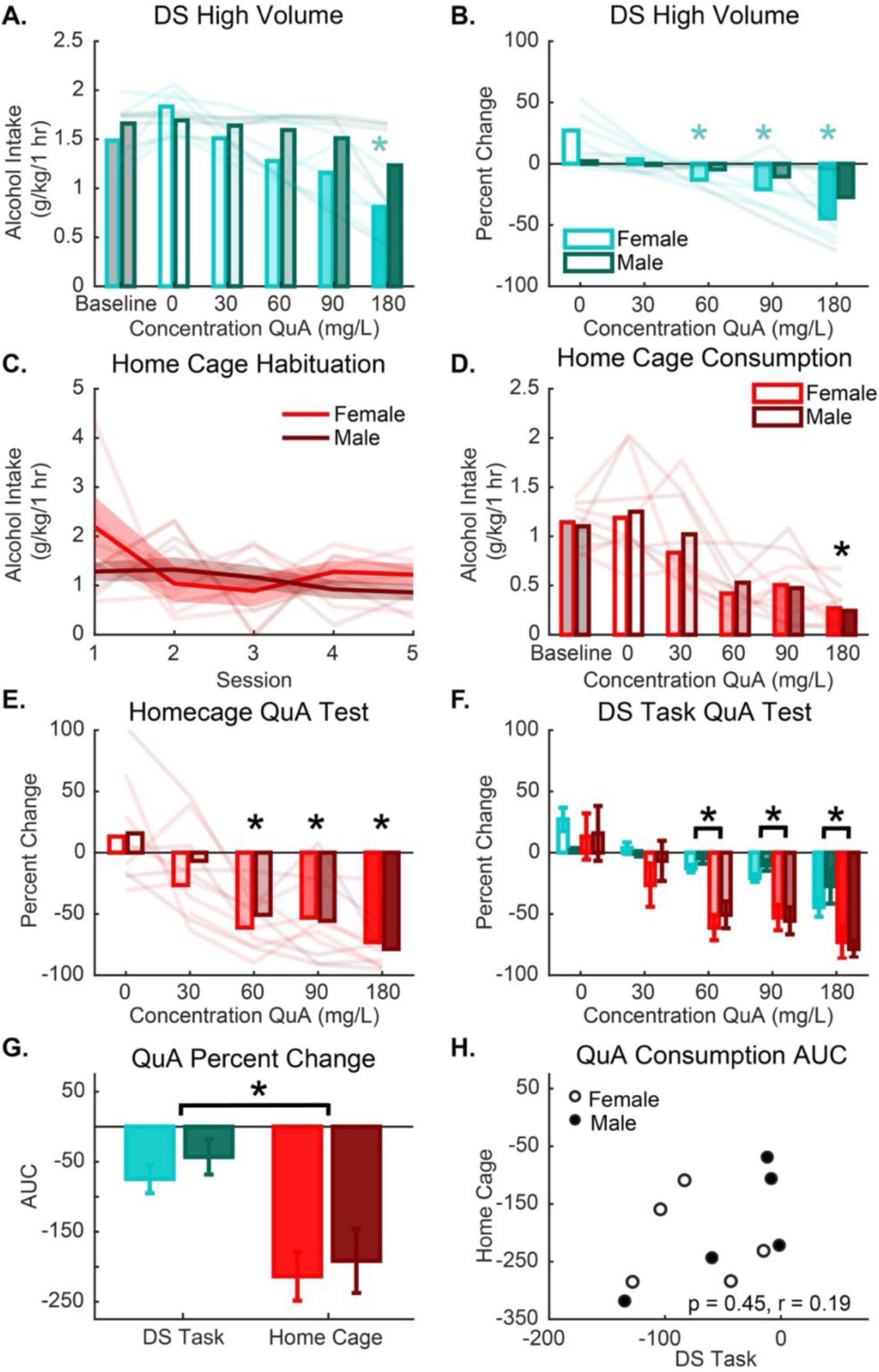
Rats demonstrate greater aversion-resistant drinking in the High-Volume DS task compared with home cage drinking. **A.** Quinine-alcohol consumption was not altered across increasing concentrations of quinine except at 180 mg/L in female rats only. **B.** Consumption as percent change from baseline demonstrated that female rats significantly reduced their consumption at 60, 90, and 180 mg/L. **C.** Rats habituated to the home cage for five sessions across five days. **D.** Quinine-alcohol consumption was reduced in the home cage at 180 mg/L for both male and female rats. **E.** Consumption as percent change from baseline was significantly reduced at 60, 90, and 180 mg/L for both male and female rats. **F.** When we assessed percent change in consumption comparing DS vs home cage consumption, we found that DS task consumption was significantly less suppressed than home cage at 60, 90, and 180 mg/L. **G.** Area under the curve (AUC) of the percent change in quinine-alcohol consumption was significantly more suppressed in home cage than in the DS task. **H.** AUC scores in the home cage were not predictive of AUC scores in the DS task. Color coded asterisks (*) indicate a significant difference within sex, black asterisks with a bar over them indicate a significant difference between the DS task and home cage, black asterisks without a bar above them indicate a significant group difference (collapsed by sex) compared with control conditions.

Next, we assessed how consumption was affected by quinine adulteration as a percent change from baseline. Quinine adulteration reduced consumption (Fig. 4B, main effect of quinine concentration in a sex-dependent manner: F(1,40) = 65.34 p < 0.0001, η² = 0.62; main effect of sex: F(1,24.66) = 3.13, p = 0.09, η² = 0.11; interaction of sex and concentration: F(1,40) = 9.54, p = 0.003, η² = 0.193). Post-hoc Bonferroni corrected Dunnet’s t-test determined that compared with 0 mg/L, all concentrations but 30 mg/L significantly reduced consumption (0 vs 30 mg/L: *diff* = −13.29, 95% CI [7.18, −33.77], p = 0.30; 0 vs 60 mg/L: *diff* = −23.42, 95% CI [−43.91, −2.96], p = 0.02; 0 vs 90 mg/L: *diff* = −30.26, 95% CI [−50.74, −9.79], p = 0.002; 0 vs 180 mg/L: *diff* = −50.55, 95% CI [−71.03, −30.07], p < 0.0001). Further post-hoc tests determined that this was driven by females as Dunnet’s t-test in females only revealed that 0 mg/L significantly differed from 60 mg/L (*diff* = −0.56, 95% CI [−1.00, −0.11], p = 0.01), 90 mg/L (*diff* = −0.67, 95% CI [−1.12, −0.23], p = 0.003), and 180 mg/L (*diff* = −1.02, 95% CI [−1.47, −0.58], p < 0.0001). Post-hoc comparison of males vs females at each concentration revealed no significant sex differences (p’s > 0.10).

We then assessed aversion-resistant drinking in the home cage to determine whether aversion-resistant drinking sensitivity differs in the absence of explicit alcohol cues. Prior to home cage aversion-resistant drinking testing, rats underwent 5 days of baseline home cage habituation where there was a main effect of day (Fig 4C, main effect of day: F(1,50) = 4.87, p = 0.03, η² = 0.09; no significant effect of sex or interaction of day and sex, (p’s > 0.10)). Post-hoc Dunnet’s t-test showed that consumption did not differ on any of the days compared with day 1 (all p’s > 0.10). We then tested the impact of quinine adulteration on drinking in the home cage. Quinine adulteration significantly reduced consumption (Fig 4D, main effect of quinine concentration: F(5, 49.33) = 17.19, p < 0.0001, η² = 0.64; no significant effect of sex or interaction of sex and quinine concentration (p’s > 0.10)). Post-hoc Dunnet’s test with Bonferroni correction revealed that baseline consumption significantly differed from 180 mg/L (*diff* = −0.55, 95% CI [−0.93, −0.19], p = 0.002). Percent change consumption from the baseline drinking was significantly reduced across concentration of quinine (Fig 4E; main effect of concentration: F(1, 39.21) = 37.47, p < 0.0001, η² = 0.49; and no significant effect of sex or interaction of concentration and sex, (p’s > 0.10). Post-hoc Dunnet’s t-test with Bonferroni correction revealed that 0 differed from 60 mg/L (*diff* = −70.49, 95% CI [−108.60, −32.39], p < 0.001), 90 mg/L (*diff* = −68.86, 95% CI [−106.96, −30.75], p < 0.001), and 180 mg/L (*diff* = 90.30, 95% CI [−128.40, −52.20], p < 0.0001), but not 30 mg/L (p = 0.14). These data demonstrate that alcohol drinking under free access conditions in the home cage remained aversion sensitive at most quinine concentrations that we tested.

Direct comparison of percent change scores in consumption in the DS task versus the home cage revealed that drinking was more aversion-resistant in the DS task in comparison to the home cage (Fig 4F, main effect of drinking context: F(1, 89.26) = 4.38, p = 0.04, η² = 0.05; main effect of quinine concentration: F(1, 89.18) = 77.96, p < 0.0001, η² = 0.47; no significant effect of sex or interaction of sex and quinine concentration (p’s > 0.10); interaction of sex and drinking context: F(1,89.26) = 3.23, p = 0.08, η² = 0.04; interaction of quinine concentration and drinking context: F(1, 89.18) = 4.9, p = 0.03, η² = 0.05; interaction of sex, quinine concentration, and drinking context: F(1, 89.18) = 3.9, p = 0.051, η² = 0.04). Post-hoc Bonferroni corrected Welch’s t-test comparing percent change in consumption from DS vs home cage at each quinine concentration revealed that aversion-resistant drinking was significantly stronger in the DS task vs the home cage at 60 mg/L (*t*(11.72) = 5.99, p < 0.0001), 90 mg/L (*t*(14.94) = 3.10, p = 0.007), and at 180 mg/L (*t*(17.55) = 3.69, p = 0.002). Finally, to further demonstrate the difference in aversion-resistant drinking between the DS task and the home cage, we calculated the area under the curve for the percent change scores for quinine-alcohol drinking finding that aversion-resistant drinking was stronger in the DS task compared with the home cage (Fig 4G; main effect of drinking context: F(1,10) = 37.55, p < 0.0001, η² = 0.79; no significant effect of sex or interaction of sex and drinking context (p’s > 0.10)). These data demonstrate that aversion-resistant drinking is greater in the DS task than the home cage suggesting that cues evoke aversion-resistant drinking.

### Individual differences in quinine sensitivity are correlated within behavioral context, but not across behavioral contexts

Because we observed individual variability in aversion-resistance in both the DS task and home cage, we next wanted to assess whether quinine sensitivity in one behavioral context was related to quinine sensitivity in the other behavioral context. First, we assessed the correlation between the area under the curve (AUC) scores for percent change in consumption on quinine sessions. We found that DS task AUCs did not predict home cage AUCs (Fig 4H; r(8) = .19, p = 0.45) suggesting that aversion-resistance in the DS task is not predictive of aversion-resistance during home cage drinking.

To further assess this relationship, we then calculated correlations between quinine adulteration percent change scores at each concentration of quinine-adulterated alcohol both within the behavioral context (DS task or home cage drinking) and between behavioral contexts. We found significant correlations between percent change within the DS task and within home cage drinking tests (Fig. 5, DS task: 60 mg/L predicts 90 mg/L (r(26) = 0.81, p = 0.004), 60 mg/L predicts 180 mg/L (r(26) = 0.83, p = 0.003), 90 mg/L predicts 180 mg/L (r(26) = 0.75, p = 0.01); Home cage drinking: 30 mg/L predicts 60 mg/L (r(26) = 0.64, p = 0.046). Importantly, the percent change in consumption in response to quinine adulteration in the DS task did not predict the percent change in consumption in the home cage (all p’s > 0.10). Together, these data suggest that the behavioral phenotype of cue-evoked aversion-resistant drinking is distinct from home cage aversion-resistant drinking.

**Figure 5.**
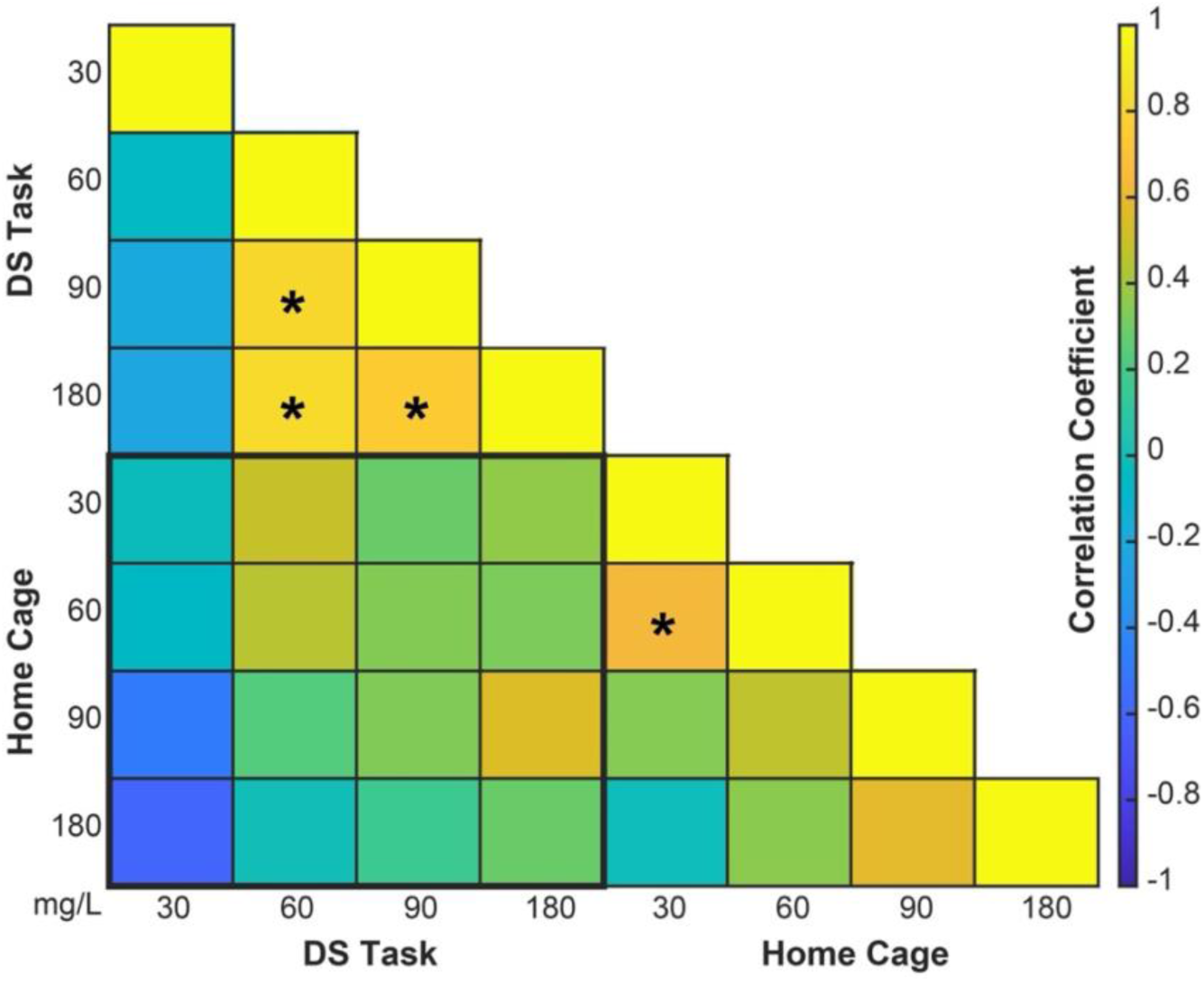
Individual differences in quinine sensitivity are correlated within behavioral context, but not across behavioral contexts. Correlation matrix analyses of the high-volume DS task and home cage revealed that DS task consumption at 60 mg/L positively predicted DS task consumption at 90 and 180 mg/L and 90 mg/L predicted 180 mg/L. 30 mg/L positively predicted home cage consumption at 60 mg/L. Importantly, consumption in the DS task was not predictive of home cage consumption at any level. Asterisks (*) indicate a significant difference and heatmap colors indicate r^2^ values.

## DISCUSSION

Here we demonstrate for the first time that alcohol cues can invigorate aversion-resistant drinking. This effect is greater than aversion-resistant drinking in environments that are free of explicit alcohol-related cues (i.e., the home cage). We found that aversion-resistant drinking in the standard version of the DS task with limited alcohol availability was robust and was not reduced by any concentration of quinine-alcohol. When we increased the reward volume so that rats could titrate their consumption levels, we found aversion-resistant drinking was still robust and only significantly reduced in female rats at 180 mg/L, the highest concentration of quinine we tested. However, when we assessed the consumption levels as percent change from averaged baseline days, we found that consumption reduced at 60, 90, and 180 mg/L in female, but not male rats. As a control for an environment free of explicit alcohol-related cues we assessed aversion-resistant drinking in the same rats in the home cage. We found that rats displayed aversion-resistant drinking at all quinine-alcohol concentrations except 180 mg/L. When we assessed aversion-resistant drinking as percent change from averaged baseline days, we found that both male and female rats significantly reduced consumption at 60, 90, and 180 mg/L. Importantly, when we compared percent change consumption levels in the DS task versus the home cage, we found that DS task quinine-alcohol drinking was significantly greater than home cage, suggesting invigorated aversion-resistant drinking in the DS task, relative to the home cage. To further support this finding, DS task and home cage quinine-alcohol consumption were not predictive of one another. Finally, we created a correlation matrix of DS task quinine-alcohol consumption with home cage quinine-alcohol consumption across all concentrations. We found that quinine-alcohol intake was predictive within, but not across, environmental context suggesting that aversion-resistant drinking in the DS task may be a construct that is unique and separable from home cage aversion-resistant drinking. In summary, we found that aversion-resistant alcohol drinking is stronger in environments with alcohol related cues. To our knowledge this is the first causal evidence that cues can invigorate aversion-resistant drinking.

Previous research has used cue-based tasks in assessing alcohol-related behaviors. To our knowledge this is the first data that shows a commonly used feature of behavior training, cues, invigorate aversion-resistant drinking to a greater extent than aversion-resistant drinking in an environment free of explicit alcohol-related cues. Cues are widely known to drive relapse and craving (Vafaie & Kober, 2022) so it isn’t particularly surprising that cues would also invigorate aversion-resistant drinking. Despite this being an intuitive finding, it is the first of its kind and maintains important implications. Previous research on aversion-resistant alcohol seeking and consumption has used behavioral models that incorporate cues (Bauer, 2023; Giuliano et al., 2021; Grodin et al., 2018; Hopf et al., 2010; McCane et al., 2021; Siciliano et al., 2019; Timme et al., 2020, 2022, 2024; Ward et al., 2025), but prior work has not directly assessed the impact of those cues. Our findings may have important implications for these types of experiments. For example, head-fixed mice responding to Pavlovian conditioned stimuli for alcohol demonstrate robust aversion-resistant drinking that is absent when these mice are tested in their home cage (Timme et al., 2024). While this finding was attributed to head fixation, our findings in freely moving rats suggest that this aversion-resistant drinking may have been more related to the presence of alcohol cues. Thus, our findings have important implications for the field and for anyone moving forward with assessment of aversion-resistant drinking in an alcohol-related cue-based task. Additionally, an important real-world direction for reducing harmful alcohol use may be implementing limits on alcohol-related cues.

The role of cues in driving aversion-resistant drinking may also explain the strong degree of overlap in the neural mechanisms for cue-evoked and aversion-resistant alcohol seeking and consumption. For example, neural circuits underlying cue-based craving include the nucleus accumbens, prefrontal cortex, and dorsal striatum (Pickens et al., 2011; Ray & Roche, 2018; Seo & Sinha, 2014). For instance, in a human laboratory setting where participants received a shock if they responded to an alcohol cue, Grodin et al. (2018) found that high drinkers had greater activity in the dorsal striatum, nucleus accumbens, and dorsomedial prefrontal cortex during trials where participants responded to an alcohol cue despite a high likelihood of being shocked. Relative to low drinkers, high drinkers also had greater functional connectivity between the insula and nucleus accumbens during high threat alcohol cues and this connectivity positively predicted compulsive-like drinking score (Grodin et al., 2018). The role of overlapping neural circuitry in cue-elicited and aversion-resistant drinking is further supported by the preclinical literature: aversion-resistant drinking depends on the nucleus accumbens (Sneddon et al., 2021), the dorsal striatum (da Silva et al., 2024; Bauer et al., 2026; Patton et al., 2021), prefrontal cortex (Seif et al., 2013; Timme et al., 2022), and insula (Fornari et al., 2026; Starski et al., 2024). Thus, a useful potential therapeutic target to combat cue-invigorated aversion-resistant drinking may be an already identified cue-related brain region.

Our findings are consistent with the incentive sensitization theory of addiction, and specifically, the prediction that cues produce motivational states that invigorate aversion-resistant drinking. Incentive sensitization posits that addiction is driven, at least in part, by the ability of drug and alcohol cues to promote enhanced ‘wanting’ even while ‘liking’ of addictive substances is decreased (Berridge & Robinson, 2016; Robinson & Berridge, 1993, 2001, 2025). Indeed, in an updated review of incentive sensitization theory, Robinson & Berridge (2025) suggest that motivational “compulsion”, as seen in continued drug use despite negative consequences may be due to excessive incentive salience. Thus, intense ‘wanting’ for alcohol in the presence of alcohol-related cues could drive continued alcohol use despite a direct experimental reduction in ‘liking’ or the palatability of an alcohol reward by the addition of quinine. Importantly, drug-related cues may be better able to induce a state of motivational compulsion or ‘wanting’. Indeed, compared with cues for natural reinforcers like sucrose, drug-related cues drive craving and relapse to a greater extent and across a longer duration (Bienkowski et al., 2004; Ciccocioppo et al., 2004; Madangopal et al., 2026; Vafaie & Kober, 2022). Thus, while cues for non-drug rewards may also elicit ‘wanting’ and compulsive-like seeking and consumption, we hypothesize that these effects are stronger for cues predicting addictive drugs and particularly relevant for modeling compulsive drug and alcohol use.

Our results add to the growing body of literature surrounding the theoretical side of aversion-resistance. Our findings suggest that general motivation may be increased by cues, though it is difficult to ascertain the effect of cues on motivation purely from our behavior and experimental design. Importantly, not all compulsive behaviors are “aversion resistant” and addiction is not solely defined by insensitivity to negative outcomes or compulsivity (George, Ahmed, & Gilpin, 2022). Others have suggested that what appears to be resistance to punishment in the human population is due to excessive goal-directed drug choice or motivation for drug (Hogarth, 2020). Whether our findings are related to more goal-directed alcohol seeking and/or general increases in motivation, and whether they extend to other models of addiction or compulsion is an important area of future research.

Despite the robust aversion-resistant alcohol drinking in our findings, we are confident that the rats still found the quinine adulteration aversive. One clue to this is that as quinine-concentration increased, NS responding decreased. We found that responding during the NS was significantly reduced at 90 and 180 mg/L for both the low- and high-volume rewards. We initially considered that this may be due to a “day of the week” effect as both 90 and 180mg/L were administered on Thursday and Friday, respectively. However, we assessed NS responding during alcohol only drinking weeks and did not see a reduction in NS responding at any concentration. Further, when we looked at alcohol only NS responding following 180 mg/L quinine-alcohol, NS responses significantly increased, further suggesting that the reduced NS response during quinine is not due to extinguished responding and may be meaningful to the quinine stimulus. These data provide further evidence that the rats find the quinine-alcohol aversive and contributes to the growing body of literature revealing several behavioral indicators of aversion-resistant drinking such as rate and pattern of consumption (Bauer et al., 2021; Darevsky et al., 2019) as well as facial reactions (Timme et al., 2024). Previous research suggests a “head-down and push” strategy whereby consumption of highly aversive quinine-concentrations evokes a constrained behavioral pattern of consumption (Darevsky et al., 2019). The reduced NS responding at high quinine-concentrations that we observe could suggest a constrained reward seeking approach similar to the head-down and push theory. Thus, our data suggest that rats confine reward seeking to the DS more so during increasingly aversive alcohol drinking which may be a behavioral strategy in aversion-resistant alcohol drinking.

Our sex-biased effects are distinct from previous findings. In prior work on aversion-resistant drinking that has used males and females, sex-related effects are mixed. While some studies have found no sex differences in aversion-resistant drinking (Bauer et al., 2021; 2026; Randall et al., 2017) studies that identify a sex effect generally report that females are more aversion resistant than males (DeBaker et al., 2020; Fornari et al., 2026; Sneddon et al., 2019). One exception in the literature is from selectively bred alcohol-preferring P rats: male P rats develop greater aversion-resistance than female P rats after chronic ethanol drinking (Katner et al., 2022). These differences could be due to differences in species, strain, alcohol exposure patterns, testing procedures or methods of calculating aversion-resistance. Here, we find no sex differences in aversion-resistant drinking in the home cage or in the low-volume version of the DS task, but that males are more aversion-resistant in the DS task when higher volumes of alcohol are available. Despite moderate sample sizes when split by sex (n = 5/sex), we report robust effect sizes for main effects and interactions of sex and quinine-alcohol for this effect. Greater aversion-resistance in males in the DS task may be due to limitations on front-loading as the DS task does not allow front-loading, which may be a necessary behavioral driver of aversion-resistant drinking in females, but not males. Alternatively, the volumes used in the high-volume version of the task may still constrain alcohol consumption at baseline in males: males continue to response to the DS almost 100% of the time, whereas females reduce their responding to the DS before stabilizing. While both males and females are able to earn and consume amounts of alcohol equal to or greater to what they choose to consume during free access in the home cage, this may still represent constrained consumption for males in the DS task, which could contribute to greater aversion-resistance.

In conclusion, we find that alcohol cues invigorate aversion-resistant drinking. Our findings have implications for the broader field of behavioral neuroscience where cues are incorporated into training and testing of aversion-resistant drinking. We provide the first behavioral model of cue-invigorated aversion-resistant drinking, which appears to be a unique construct when compared with aversion-resistant drinking in environments free of explicit reward-paired cues as home cage and DS task aversion-resistant drinking were not predictive of one another. We also found that female rats are more sensitive to quinine adulteration of alcohol than males in some conditions when they have undergone DS task training and DS task quinine-alcohol testing. Future research will utilize this model when assessing the neurobiological mechanism of cue invigorated aversion-resistant drinking.

