## Supplemental Tables and Figure for "Alcohol Cues Invigorate Aversion-Resistant Alcohol Drinking"

Supplementary Material

**Supplementary Table 1. Non-significant results from omnibus testing**

| **Figure** |  | **Degrees of Freedom** | **F-value** | **P-value** | **Effect Size (η²)** |
| --- | --- | --- | --- | --- | --- |
| **1B** | **Alcohol Consumption:** Day X Sex | | | | |
|  | Main effect of day | 1, 110 | 1.83 | 0.18 | 0.02 |
|  | Main effect of sex | 1, 14.3 | 2.79 | 0.12 | 0.16 |
|  | Interaction of sex and day | 1, 110 | 2.22 | 0.14 | 0.02 |
| **1C** | **DS Port Entries**: Day X Sex | | | | |
|  | Main effect of sex | 1, 11.85 | 2.11 | 0.17 | 0.15 |
|  | Interaction of sex and day | 1, 110 | 1.18 | 0.28 | 0.01 |
|  | **Port Entries**: Day X Sex X Cue Type | | | | |
|  | Interaction between cue type, sex, and day | 1,150 | 0 | 0.95 | 0 |
| **1D** | **Number of Rewards:** Day X Sex |  |  |  |  |
|  | Main effect of sex | 1, 11.85 | 2.1 | 0.17 | 0.15 |
|  | Interaction of sex and day | 1,210 | 1.16 | 0.2836 | 0.005 |
| **1E** | **Alcohol Consumption:** Day X Sex | | | | |
|  | Main effect of sex | 1,12.99 | 0.01 | 0.923 | 0.001 |
| **2A** | **DS responses:** Sex X Quinine Concentration | | | | |
|  | Main effect of quinine adulteration | 5,50 | 1.33 | 0.27 | 0.12 |
|  | Main effect of sex | 1,10 | 1.62 | 0.23 | 0.14 |
|  | Interaction of sex and quinine concentration | 5,50 | 0.75 | 0.59 | 0.07 |
| **2B** | **Percent Change in DS Responses:** Sex X Quinine Concentration | | | | |
|  | Main effect of quinine adulteration | 1,40 | 0.2 | 0.6 | 0.01 |
|  | Main effect of sex | 1,18.92 | 0.18 | 0.68 | 0.01 |
|  | Interaction of sex and quinine concentration | 1,40 | 1.92 | 0.17 | 0.05 |
| **2C** | **NS Responses:** Sex X Quinine Concentration | | | | |
|  | Main effect of sex | 1,10 | 0.02 | 0.90 | 0.002 |
|  | Interaction of sex and quinine concentration | 5,50 | 1.38 | 0.25 | 0.121 |
| **2D** | **Percent Change in NS Responses:** Sex X Quinine Concentration | | | | |
|  | Main effect of sex | 1,50 | 0.01 | 0.75 | 0.002 |
|  | Interaction of sex and quinine concentration | 1,50 | 0.69 | 0.41 | 0.014 |
| **1C** | **NS Responding M-F, across training days:** Day/Quinine Concentration X Sex | | | | |
|  | Main effect of sex | 1,50 | 0.41 | 0.52 | 0.01 |
|  | Interaction of sex and day/quinine concentration | 1,40 | 1.17 | 0.29 | 0.028 |
| **2E** | **DS Port Entry Latency:** Quinine Concentration X Sex | | | | |
|  | Main effect of sex | 1,10 | 0.69 | 0.42 | 0.065 |
|  | Main effect of quinine concentration | 5, 50 | 1.71 | 0.15 | 0.146 |
|  | Interaction of sex and quinine concentration | 5, 50 | 0.51 | 0.76 | 0.049 |
| **2F** | **NS Port Entry Latency:** Quinine Concentration X Sex | | | | |
|  | Main effect of sex | 1, 10 | 2.94 | 0.117 | 0.227 |
|  | Main effect of quinine concentration | 5, 50 | 0.76 | 0.5821 | 0.071 |
| **2G** | **Quinine Consumption:** Sex X Quinine Concentration | | | | |
|  | Main effect of quinine concentration | 5,50 | 0.68 | 0.64 | 0.06 |
|  | Interaction of sex and quinine concentration | 5,50 | 0.54 | 0.74 | 0.05 |
| **2H** | **Percent Change in Quinine Consumption:** Sex X Quinine Concentration | | | | |
|  | Main effect of quinine concentration | 1, 50 | 1.33 | 0.25 | 0.03 |
|  | Main effect of sex | 1, 50 | 2.12 | 0.1515 | 0.04 |
|  | Interaction of sex and concentration | 1, 50 | 2.33 | 0.13 | 0.05 |
| **3A** | **Port Entry Probability:** Sex X Cue Type X Day | | | | |
|  | Main effect of sex | 1, 25.18 | 2.36 | 0.14 | 0.09 |
| **3B** | **Alcohol Consumption Days 6 - 9:** Sex X Cue Type X Day | | | | |
|  | Main effect of sex | 1, 39.31 | 1.86 | 0.18 | 0.05 |
|  | Interaction of sex and day | 1, 30 | 0.49 | 0.49 | 0.02 |
| **3D** | **Percent Change in DS Responses:** Sex X Quinine Concentration | | | | |
|  | Main effect of sex | 1, 10 | 0.38 | 0.55 | 0.037 |
| **3E** | **NS Responses:** Sex X Quinine Concentration | | | | |
|  | Main effect of sex | 1, 10 | 2.11 | 0.18 | 0.17 |
| **3F** | **Percent Change in NS Responses:** Sex X Quinine Concentration | | | | |
|  | Interaction of sex and quinine concentration | 4, 40 | 0.70 | 0.60 | 0.07 |
| **3H** | **Latency to Port Entry Following NS:** Sex X Quinine Concentration | | | | |
|  | Main effect of sex | 1, 57 | 2.18 | 0.145 | 0.037 |
|  | Main effect of quinine concentration | 5, 57 | 0.77 | 0.5752 | 0.063 |
|  | Interaction of sex and quinine concentration | 5, 57 | 1.93 | 0.104 | 0.145 |
| **4A** | **Quinine Alcohol Consumption:** Sex X Quinine Concentration | | | | |
|  | Main effect of sex | 1, 10 | 2.33 | 0.16 | 0.19 |
| **4C** | **Home Cage Consumption:** Sex X Day | | | | |
|  | Main effect of sex | 1, 50 | 0.60 | 0.44 | 0.01 |
|  | Interaction of sex and day | 1, 50 | 0.11 | 0.74 | 0.0 |
| **4D** | **Home Cage Quinine Alcohol Consumption:** Sex X Quinine Alcohol Concentration | | | | |
|  | Main effect of sex | 1, 10.19 | 0.15 | 0.70 | 0.02 |
|  | Interaction of sex and quinine concentration | 5, 49.33 | 0.25 | 0.94 | 0.03 |
| **4E** | **Percent Change in Home Cage Quinine Alcohol Consumption:** Sex X Quinine Alcohol Concentration | | | | |
|  | Main effect of sex | 1, 26.78 | 0.85 | 0.37 | 0.03 |
|  | Interaction of sex and quinine concentration | 1, 39.21 | 0.69 | 0.41 | 0.02 |
| **4F** | **Percent Change in Quinine Alcohol Consumption:** Drinking Context X Quinine Concentration X Sex | | | | |
|  | Main effect of sex | 1, 24.95 | 0 | 0.97 | 0.0 |
|  | Interaction of sex and quinine concentration | 1, 89.18 | 0.31 | 0.58 | 0.0 |
| **4G** | **Area Under the Curve for Percent Change Scores in Quinine-Alcohol Drinking:** Drinking Context X Sex | | | | |
|  | Main effect of sex | 1, 10 | 0.6 | 0.46 | 0.06 |
|  | Interaction of sex and drinking context | 1, 10 | 0.04 | 0.85 | 0.004 |
| **Sup. 1A** | **Percent Change in Latency, Low Volume DS Responding:** Sex X Quinine Concentration | | | | |
|  | Main effect of sex | 1, 50 | 0.07 | 0.7955 | 0.026 |
|  | Main effect of quinine concentration | 4, 50 | 0.33 | 0.8577 | 0.026 |
|  | Interaction of sex and quinine concentration | 4, 50 | 1.04 | 0.3936 | 0.077 |
| **Sup. 1B** | **Percent Change in Latency, Low Volume NS Responding:** Sex X Quinine Concentration | | | | |
|  | Main effect of sex | 1, 10 | 1.41 | 0.2618 | 0.124 |
|  | Main effect of quinine concentration | 4, 40 | 1.12 | 0.3592 | 0.101 |
|  | Interaction of sex and quinine concentration | 4, 40 | 2.11 | 0.098 | 0.174 |
| **Sup. 1C** | **Percent Change in Latency, High Volume DS Responding:** Sex X Quinine Concentration | | | | |
|  | Main effect of sex | 1, 10 | 1.06 | 0.3265 | 0.096 |
|  | Interaction of sex and quinine concentration | 4, 40 | 2.45 | 0.062 | 0.197 |
| **Sup. 1D** | **Percent Change in Latency, High Volume NS Responding:** Sex X Quinine Concentration | | | | |
|  | Main effect of sex | 1, 10 | 0.03 | 0.8664 | 0.003 |
|  | Main effect of quinine concentration | 4, 40 | 0.97 | 0.4367 | 0.088 |
|  | Interaction of sex and quinine concentration | 4, 40 | 2.1 | 0.0993 | 0.173 |

**Supplementary Table 2.** **Non-significant results from post-hoc pair-wise comparisons**

| **Figure** | **Dunnet’s Multiple Comparisons Test** | **Diff** | **Confidence Interval [lower, upper]** | **P-value** |
| --- | --- | --- | --- | --- |
| **2C** | **NS Port Entry Probability:** Baseline vs all other quinine concentrations | | | |
|  | Baseline vs 0 mg/L | -0.0756 | [-0.2298, 0.0786] | 0.58579 |
|  | Baseline vs 30 mg/L | 0.0114 | [-0.1428, 0.1656] | 0.99981 |
|  | Baseline vs 60 mg/L | 0.0024 | [-0.1518, 0.1566] | 1.00000 |
| **2D** | **NS Port Entry Probability Percent Change:** Zero vs all other quinine concentrations | | | |
|  | 0 vs 30 mg/L | 19.9484 | [-9.8263, 49.7232] | 0.2757 |
|  | 0 vs 60 mg/L | 21.85943 | [-7.9153, 51.6342] | 0.2066 |
| **1C** | **NS Port Entry Probability:** Monday vs all other days of the week | | | |
|  | Monday vs Tuesday | -0.060 | [-0.2216, 0.1016] | 0.7531 |
|  | Monday vs Wednesday | 0.044 | [-0.1176, 0.2056] | 0.8968 |
|  | Monday vs Thursday | 0.017 | [-0.1446, 0.1786] | 0.9965 |
|  | Monday vs Friday | 0.065 | [-0.0966, 0.2266] | 0.7002 |
| **2F** | **Latency to Enter Port Following NS Low Volume:** Females – Baseline vs all other quinine concentrations | | | |
|  | Baseline vs 0 mg/L | -0.16833253 | [-2.3616, 2.0249] | 0.9997 |
|  | Baseline vs 30 mg/L | -0.03420077 | [-2.2274, 2.1590] | 1.0000 |
|  | Baseline vs 60 mg/L | 0.13752672 | [-2.0557, 2.3308] | 0.9999 |
|  | Baseline vs 90 mg/L | 0.61109418 | [-1.5821, 2.8043] | 0.3041 |
|  | Baseline vs 180 mg/L | 1.42121481 | [-0.7720, 3.6144] | 0.9095 |
|  | **Latency to Enter Port Following NS Low Volume:** Males – Baseline vs all other quinine concentrations | | | |
|  | Baseline vs 0 mg/L | 0.21399022 | [-1.0964, 1.5244] | 0.9895 |
|  | Baseline vs 30 mg/L | 0.04662659 | [-1.2638, 1.3571] | 1.000 |
|  | Baseline vs 60 mg/L | 0.10918649 | [-1.2012, 1.4196] |  |
|  | Baseline vs 90 mg/L | -0.37311530 | [-1.6835, 0.9373] | 0.9024 |
|  | Baseline vs 180 mg/L | -0.29625974 | [-1.6067, 1.0142] | 0.9588 |
| **3C** | **DS Port Entry Probability Across Quinine Concentrations, High Volume:** Females – Baseline vs all other quinine concentrations | | | |
|  | Baseline vs 30 mg/L | -0.0108 | [-0.1602, 0.1386] | 0.9998 |
|  | Baseline vs 60 mg/L | -0.0828 | [-0.2322, 0.0667] | 0.4446 |
|  | Baseline vs 90 mg/L | -0.1088 | [-0.2582, 0.0406] | 0.2115 |
|  | **DS Port Entry Probability Across Quinine Concentrations, High Volume:** Males – Baseline vs all other quinine concentrations | | | |
|  | Baseline vs 0 mg/L | 0.0224 | [–0.1722, 0.2170] | 0.9979 |
|  | Baseline vs 30 mg/L | 0.0024 | [-0.1922, 0.1970] | 1.0000 |
|  | Baseline vs 60 mg/L | -0.0296 | [-0.2242, 0.1650] | 0.9923 |
|  | Baseline vs 90 mg/L | -0.0496 | [-0.2442, 0.1450] | 0.9346 |
|  | Baseline vs 180 mg/L | -0.0636 | [-0.2582, 0.1310] | 0.8455 |
| **3D** | **Percent Change in DS Port Entry Probability:** Males – 0 mg/L vs all other quinine concentrations | | | |
|  | 0 vs 30 mg/L | -2.068966 | [-18.1288, 13.9909] | 0.9906 |
|  | 0 vs 60 mg/L | -6.074271 | [-22.1341, 9.9856] | 0.7128 |
|  | 0 vs 90 mg/L | -8.381963 | [-24.4418, 7.6779] | 0.4609 |
|  | 0 vs 180 mg/L | -9.735632 | [-25.7955, 6.3242] | 0.3349 |
| **3G** | **Latency to Port Entry Following DS:** Females - Baseline vs all other quinine concentrations | | | |
|  | Baseline vs 0 mg/L | -0.7714 | [-0.9111, 0.8615] | 0.1031 |
|  | Baseline vs 30 mg/L | -0.0765 | [-0.9628, 0.8098] | 0.9995 |
|  | Baseline vs 60 mg/L | -0.1187 | [-1.005, 0.7676] | 0.9957 |
|  | Baseline vs 90 mg/L | -0.0248 | [-0.9111, 0.8615] | 1.0000 |
|  | Baseline vs 180 mg/L | 0.2343 | [-0.6520, 1.1206] | 0.9251 |
|  | **Latency to Port Entry Following DS:** Males - Baseline vs all other quinine concentrations | | | |
|  | Baseline vs 0 mg/L | 0.3379424 | [-0.6057, 1.2816] | 0.7577 |
|  | Baseline vs 30 mg/L | 0.1507470 | [-0.7929, 1.0944] | 0.9863 |
|  | Baseline vs 60 mg/L | 0.4173794 | [-0.5263, 1.3610] | 0.6037 |
|  | Baseline vs 90 mg/L | 0.4467796 | [-0.4969, 1.3904] | 0.5470 |
|  | Baseline vs 180 mg/L | 0.5054996 | [-0.4382, 1.4492] | 0.4417 |
| **4A** | **Quinine Alcohol Consumption High Volume:** Female - Baseline vs all other quinine concentrations | | | |
|  | Baseline vs 0 mg/L | 0.34821918 | [-0.1383, 0.8347] | 0.2241 |
|  | Baseline vs 30 mg/L | 0.02541137 | [-0.4611, 0.5119] | 1.0000 |
|  | Baseline vs 60 mg/L | 0.20691285 | [-0.6934, 0.2796] | 0.6765 |
|  | Baseline vs 90 mg/L | 0.32642442 | [-0.8129, 0.1601] | 0.2752 |
| **4A** | **Quinine Alcohol Consumption High Volume:** Male - Baseline vs all other quinine concentrations | | | |
|  | Baseline vs 0 mg/L | 0.03244350 | [-0.5248, 0.5897] | 0.9999 |
|  | Baseline vs 30 mg/L | -0.02046217 | [-0.5777, 0.5368] | 1.0000 |
|  | Baseline vs 60 mg/L | -0.06572771 | [-0.6230, 0.4915] | 0.9976 |
|  | Baseline vs 90 mg/L | -0.14961863 | [-0.7069, 0.4076] | 0.9209 |
|  | Baseline vs 180 mg/L | -0.42796295 | [-0.9852, 0.1293] | 0.1744 |
| **4B** | **Percent Change in Quinine Alcohol Consumption High Volume:** Male – Zero versus all other quinine concentrations | | | |
|  | 0 vs 30 mg/L | -0.0529 | [-0.6434, 0.5376] | 0.9976 |
|  | 0 vs 60 mg/L | -0.0981 | [-0.6887, 0.4924] | 0.9762 |
|  | 0 vs 90 mg/L | -0.1820 | [-0.7726, 0.4085] | 0.8290 |
|  | 0 vs 180 mg/L | -0.4604 | [-1.0509, 0.1301] | 0.1556 |
| **4D** | **Home Cage Quinine Alcohol Consumption:** Baseline vs All Other Quinine Concentrations | | | |
|  | Baseline vs 0 mg/L | 0.1903 | [-0.1859, 0.5665] | 0.5568 |
|  | Baseline vs 30 mg/L | 0.0024 | [-0.3737, 0.3787] | 1.0000 |
|  | Baseline vs 60 mg/L | -0.1363 | [-0.5125, 0.2399] | 0.8129 |
|  | Baseline vs 90 mg/L | -0.2380 | [-0.6142, 0.1382] | 0.3463 |
| **Figure** | **Dunn’s Test** | | **z-statistic** | **p-value** |
| **2D** | **Percent Change in NS responding:** 0 mg/l vs all other concentrations | | | |
|  | 0 vs 30 mg/L | | -14.50 | 0.2569 |
|  | 0 vs 60 mg/L | | -14.55 | 0.2519 |
| **Figure** | **Students t-test** | **t-statistic** | **Degrees of Freedom** | **p-value** |
| **2F** | **Latency to Enter Port Following NS Low Volume:** Male vs Female at every concentration of quinine | | | |
|  | M vs F: Baseline | 0.1062 | 7.8733 | 0.0918 |
|  | M vs F: 0 mg/L | 0.1449 | 7.6164 | 0.8885 |
|  | M vs F: 30 mg/L | 0.4567 | 6.0392 | 0.6638 |
|  | M vs F: 60 mg/L | 0.5915 | 5.1615 | 0.5791 |
|  | M vs F: 90 mg/L | 1.6535 | 7.9987 | 0.1368 |
|  | M vs F: 180 mg/L | 2.0086 | 4.5528 | 0.1064 |
| **4A** | **Quinine Alcohol Consumption High Volume:** Male vs Female at each Quinine Concentration | | | |
|  | M vs F: Baseline | -0.99192 | 5.2348 | 0.3648 |
|  | M vs F: 0 mg/L | 1.2891 | 5.684 | 0.2473 |
|  | M vs F: 30 mg/L | -1.1834 | 5.2507 | 0.2875 |
|  | M vs F: 60 mg/L | -1.8683 | 7.9474 | 0.09891 |
|  | M vs F: 90 mg/L | -1.4306 | 7.5268 | 0.1927 |
|  | M vs F: 180 mg/L | -1.4798 | 5.7098 | 0.1919 |
| **4B** | **Percent Change in Quinine Alcohol Consumption High Volume:** Male vs Female at each Quinine Concentration | | | |
|  | M vs F: 0 mg/L | 1.2891 | 5.684 | 0.2473 |
|  | M vs F: 30 mg/L | -1.1834 | 5.2507 | 0.2875 |
|  | M vs F: 90 mg/L | -1.4306 | 7.5268 | 0.1927 |
|  | M vs F: 180 mg/L | -1.4798 | 5.7098 | 0.1919 |

**Supplementary Table 3. Full Statistical Results from Figure 5 Correlation Matrix**

| **Correlation** | **Correlation Coefficient (r)** | **p-value** | **95% CI** |
| --- | --- | --- | --- |
| 30 DS – 30 DS | 1.0000 | - | - |
| 60 DS – 30 DS | -0.0467 | 0.89 | [-0.6570, 0.6006] |
| 90 DS – 30 DS | -0.1805 | 0. 6178 | [ -0.7275, 0.5067] |
| 180 DS – 30 DS | -0.2248 | 0.5323 | [-0.7485, 0.4715] |
| 30 HC – 30 DS | -0.0217 | 0.9526 | [-0.6425, 0.6164] |
| 60 HC – 30 DS | -0.0596 | 0.8701 | [-0.6643, 0.5922] |
| 90 HC – 30 DS | -0.4900 | 0.1505 | [-0.8557, 0.2019] |
| 180 HC – 30 DS | -0.5952 | 0.0695 | [-0.8909, 0.0550] |
| 60 DS – 60 DS | 1.0000 | - | - |
| 90 DS – 60 DS | 0.8135 | 0.0042 | [0.3770, 0.9543] |
| 180 DS – 60 DS | 0.8332 | 0.0028 | [0.4282, 0.9595] |
| 30 HC – 60 DS | 0.4981 | 0.1429 | [-0.1916, 0.8585] |
| 60 HC – 60 DS | 0.4552 | 0.1862 | [-0.2445, 0.8432] |
| 90 HC – 60 DS | 0.2323 | 0.5185 | [-0.4654, 0.7519] |
| 180 HC – 60 DS | 0.0059 | 0.9870 | [-0.6260 ,0.6332] |
| 90 DS – 90 DS | 1.0000 | - | - |
| 180 DS – 90 DS | 0.7542 | 0.0117 | [0.2372, 0.9383] |
| 30 HC – 90 DS | 0.2871 | 0.4213 | [-0.4181, 0.7764] |
| 60 HC – 90 DS | 0.3424 | 0.3328 | [-0.3662, 0.7996] |
| 90 HC – 90 DS | 0.3338 | 0.3459 | [-0.3745, 0.7961] |
| 180 HC – 90 DS | 0.1660 | 0.6467 | [-0.5177, 0.7203] |
| 180 DS – 180 DS | 1.0000 | - | - |
| 30 HC – 180 DS | 0.3726 | 0.2889 | [-0.3358, 0.8118] |
| 60 HC – 180 DS | 0.3259 | 0.3581 | [-0.3821, 0.7928] |
| 90 HC – 180 DS | 0.5489 | 0.1003 | [-0.1233, 0.8758] |
| 180 HC – 180 DS | 0.2875 | 0.4205 | [-0.4177, 0.7766] |
| 30 HC – 30 HC | 1.0000 | - | - |
| 60 HC – 30 HC | 0.6387 | 0.0468 | [-0.3631, 0.9046] |
| 90 HC – 30 HC | 0.3455 | 0.3282 | [-0.3631, 0.8009] |
| 180 HC – 30 HC | -0.0066 | 0.9856 | [-0.6336, 0.6256] |
| 60 HC – 60 HC | 1.0000 | - | - |
| 60 HC – 90 HC | 0.4762 | 0.1641 | [-0.2191, 0.8508] |
| 60 HC – 180 HC | 0.3532 | 0.3167 | [-0.3554, 0.8040] |
| 90 HC – 90 HC | 1.0000 | - | - |
| 90 HC – 180 HC | 0.5761 | 0.0813 | [-0.0840, 0.8848] |
| 180 HC – 180 HC | 1.0000 | - | - |

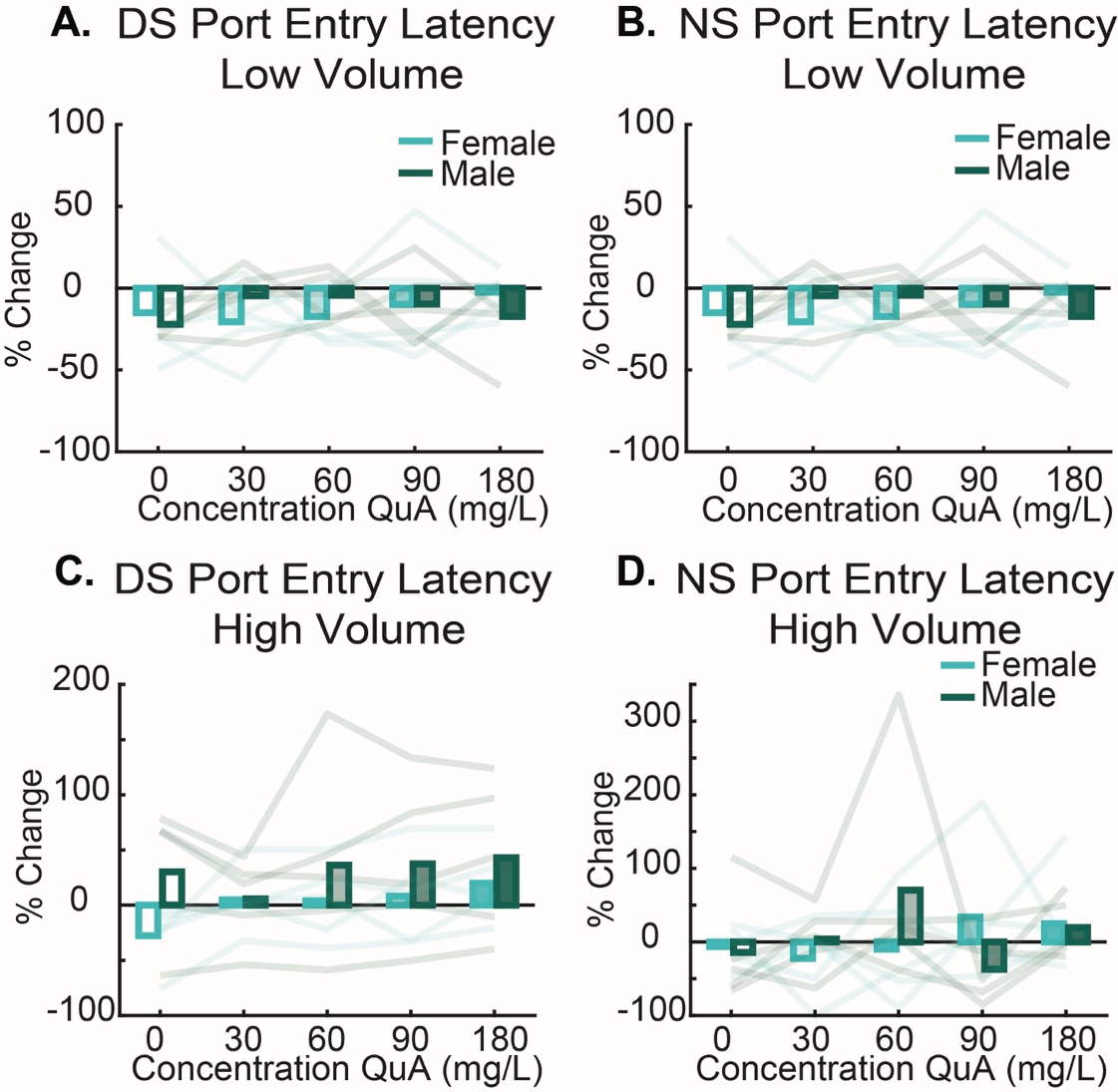

**Supplementary Figure 1. Percent Change in Latency to Respond to the Cue.** DS **(A)** nor NS **(B)** latency was affected by sex, quinine concentration, or interaction of sex and quinine concentration (p’s > 0.05). **C.** Quinine concentration significantly increased latency to respond for the DS in the high volume, F(4, 40) = 3.7, p = 0.0118, η² = 0.27, but was not altered by sex and did not interact by sex or quinine concentration (p’s > .05). **D.** NS port entry latency was unaffected by sex, quinine concentration, or interaction of sex and quinine concentration (p’s > 0.05).
